# Persistent pervasive transcription in RNA exosome depleted oocytes results in loss of female fertility

**DOI:** 10.1101/2022.04.02.486818

**Authors:** Di Wu, Jurrien Dean

## Abstract

Maturing mammalian oocytes are transcriptionally inactive and attendant RNA degradation determines the maternal transcriptome for embryonic development^1^. Perturbing oocyte RNA degradation can result in failure of meiosis, fertilization, or zygotic gene activation^1-5^. We recently reported that conditional depletion of EXOSC10, an RNA exosome associated RNase, blocks oocyte growth-to-maturation transition by interfering with ribosomal RNA processing and meiotic checkpoint genes^3^. Here we have established oocyte-specific knockout mice of a second RNA exosome associated RNase, *Dis3*. Mutant females (*Dis3*^*cKO*^) exhibit significantly reduced fertility because oocytes arrest at early maturation. DIS3 depletion allows persistent pervasive transcription, which blocks transcription termination and sequesters RNA polymerase II in intergenic regions. In addition, *Dis3*^*cKO*^ oocytes gain H3K27me3 at pre-defined loci^6^ due to insufficient demethylases KDM6A/B. Oocyte double knockout of *Dis3* and *Exosc10* causes much earlier growth defects for similar persistence of pervasive transcription, suggesting the RNA exosome complex plays a critical role to ensure transcriptome integrity during oocyte development.

---

The RNA exosome complex is responsible for most cellular RNA degradation^7^. Binding to distinct RNA exosome associated RNases, EXOSC10, DIS3 and DIS3-like, enables the complex to have different subcellular localizations and substrates^8-10^. In the absence of transcription, maturing oocytes employ RNA degradation as the driving force to shape the maternal transcriptome to support early development^1,3^. Accumulation of unwanted RNA results in extensive pathological conditions affecting oocyte maturation, fertilization, and early embryogenesis^2-4^. Discovering novel transcripts participating in chromatin remodeling, transcription termination and meiotic resumption sheds light on oocyte quality control.

To construct tissue specific *Dis3* knockout mice, exons 3-5 of the *Dis3* genomic region were flanked by loxPs using CRISPR/Cas9 (Fig. 1a). Mice with the floxed allele were crossed with *Zp3-cre* transgenic mice to conditionally disrupt *Dis3* during oocyte growth (*Dis3*^*cKO*^; Fig. 1h; Extended Data Fig. 1a-c, 3d). During 6-month harem breeding, the *Dis3*^*cKO*^ females exhibited substantially reduced fertility (Fig. 1b). There were no obvious defects in histology or weight at 12 weeks of *Dis3*^*cKO*^ ovaries which suggested successful oocyte growth (Extended Data Fig. 1d). Fully grown *Dis3*^*cKO*^ oocytes (germinal vesicle intact or GV oocytes) had normal morphology, endomembrane trafficking, and protein translation (Extended Data Fig. 1e, g). The nuclear configuration transiting from NSN (non-surrounded nucleolus) to SN (surrounded nucleolus) was intact (Fig. 1f) in contrast to the NSN dominance in *Exosc10*^*cKO*^ mice^3^. However, *ex vivo* culture of *Dis3*^*cKO*^ GV oocytes had disrupted geminal vesicle breakdown (GVBD) which could not be improved by extended culture (Fig. 1c-d; Extended Data Fig. 1h). Accordingly, we observed persistence of inhibitory phosphorylated CDK1 (pCDK1) and lack of perinuclear α-tubulin (Fig. 1e; Extended Data Fig. 1f), suggesting abrogated meiotic progression. The ovulated eggs of *Dis3*^*cKO*^ females were many fewer and composed of more immature oocytes (Extended Data Fig. 1i). In sum, *Dis3*^*cKO*^ oocytes failed to resume meiosis and arrested at the GV stage.

**Figure 1.**
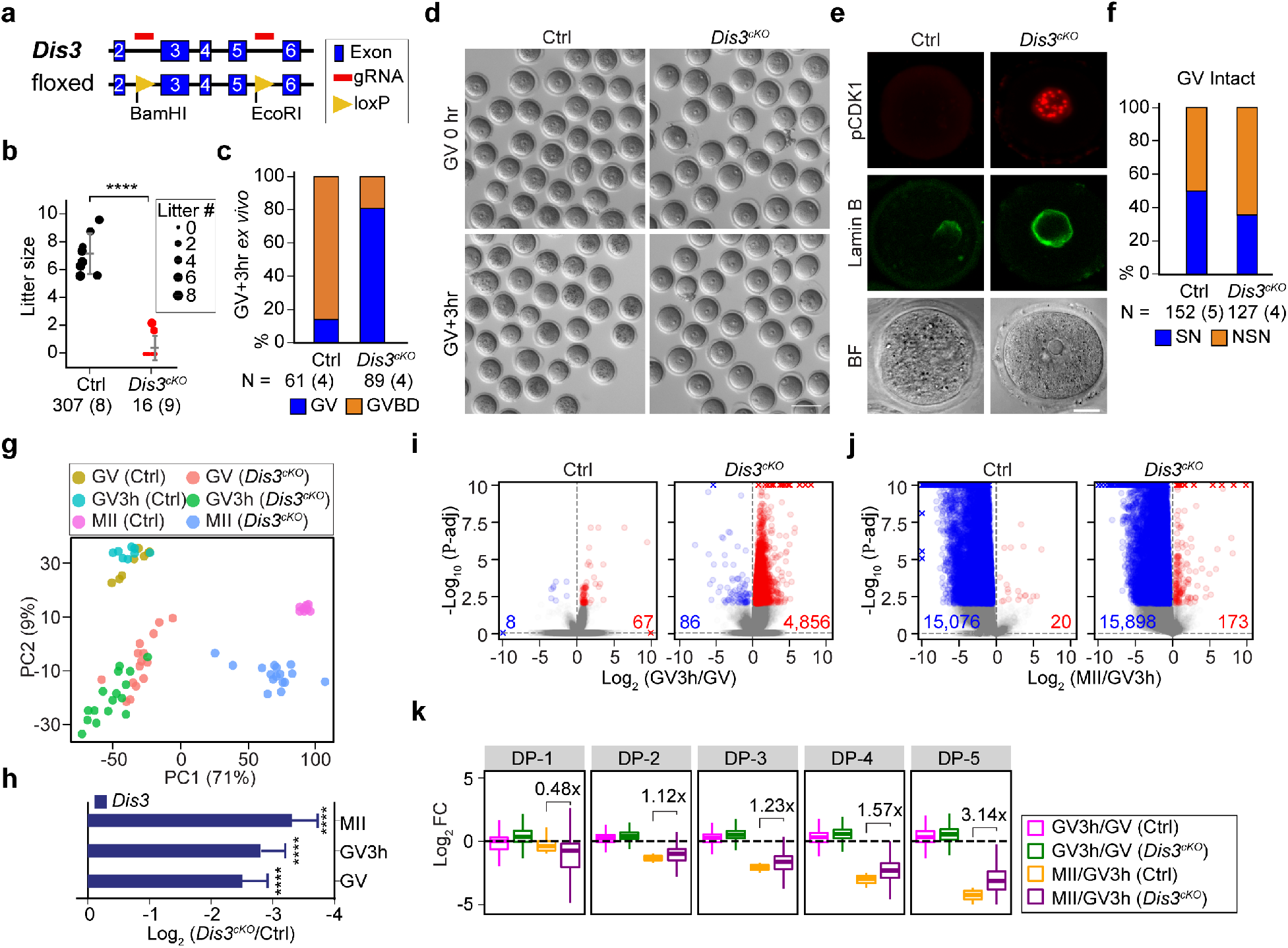
Oocyte-depletion of DIS3 causes significantly reduced female fertility due to progressive transcriptome defects during meiotic maturation. (**a**) Schematic of generating a *Dis3* floxed allele using CRISPR/Cas9. Two loxP sites were inserted surrounding exons 3-5. (**b**) Dot plot of mouse fertility. Each dot represents one female from a harem breeding of controls (Ctrl) and *Dis3*^*cKO*^ females with wildtype males. Dot sizes represent the average litter number. The number of pups and females of each genotype is indicated below each group. The horizontal lines represent the mean and standard deviation. **** P *=* 5.52e-09. (**c, d**) Percentage of GVBD oocytes after 3 hr *ex vivo* culture. The number of oocytes and females is indicated below each group. (**e**) Average intensity projection of confocal fluorescence and bright field (BF) images of lamin B and pCDK1 immunostaining at GV3h. (**f**) Bar graph showing the percentage of NSN and SN oocytes at GV stage. The number of oocytes and females is indicated below each group. (**g**) Dot plot showing principal component analysis (PCA) of all RNA-seq samples. (**h**) Bar graph showing *Dis3* transcript level decreases in *Dis3*^*cKO*^ oocytes. **** P*-*adj < 0.0001. (**i, j**) Volcano plots showing differentially expressed genes from GV to GV3h (i) and from GV3h to MII (j) in Ctrl and *Dis3*^*cKO*^ oocytes. The more and less abundant transcripts are labeled in red and blue, respectively (P-adj <0.01). (**k**) Boxplots showing the change of transcript abundance of GV3h/GV and MII/GV3h in Ctrl and *Dis3*^*cKO*^ oocytes. Degradation potential (DP) of each gene is defined by the log_2_ fold change (log_2_ FC) of MII/GV3h in Ctrl oocytes. All genes are sorted by DP in descending order and equally binned into 5 groups (DP1 to DP5). The number labeled between Ctrl and *Dis3*^*cKO*^ in each DP group represents the ratio of the mean of MII/GV3h values in *Dis3*^*cKO*^ vs. the mean of MII/GV3h values in Ctrl. Scale bars: 100 μm in d, 20 μm in e.

DIS3 disruption causes drastic changes in transcriptomes^11-13^. To visualize transcriptome dysregulation in oocytes, we performed single-oocyte RNA-seq with exogenous ERCC spike-in to facilitate inter-stage comparisons (Fig. 1g; Extended Data Fig. 2a-b; Supplementary Table 1). In both early maturation (from GV to GV3h) and mid-late maturation (from GV3h to MII), *Dis3*^*cKO*^ oocytes accumulated more transcripts than controls (Ctrl) (Fig. 1i-j). The progressive increases in transcript abundance are consistent with an overall deficiency in RNA degradation (Extended Data Fig. 2c-f). To better define the bias of DIS3 depletion in impairing RNA degradation, we defined the Degradation Potential (DP) for all transcripts according to the reduction of their abundance from GV3h to MII in wildtype oocytes. Transcripts having higher DP (DP-5) are degraded more during meiotic maturation and those having lower DP (DP-1) are degraded less (Fig. 1k). All DP groups exhibited similar transcript composition (Extended Data Fig. 2g). Interestingly, transcripts with a higher DP had a greater increase in *Dis3*^*cKO*^ than transcripts with a lower DP (Fig. 1k). A similar trend could also be observed in *Exosc10*^*cKO*^ oocytes though to a lesser extent (Extended Data Fig. 2h). We speculate that transcripts of lower DP have more protection against robust degradation, while transcripts of higher DP are less protected and prone to RNA exosome degradation^14^. In summary, *Dis3*^*cKO*^ oocytes had global and progressive RNA accumulation due to insufficient degradation.

The RNA exosome complex is known to degrade most non-coding RNA and pervasive intergenic transcripts^12^. Thus, we profiled the transcriptome at the earliest phenotype stage using unbiased RiboMinus sequencing to capture non-polyadenylated transcripts (Extended Data Fig. 3a-c; Supplementary Table 2). As expected, intergenic transcripts exhibited increased abundance in *Dis3*^*cKO*^ oocytes (Fig. 2a). PROMPTs (PROMoter uPstream Transcripts) are important intergenic transcripts and known direct targets of DIS3^11,12^. Both sense and antisense PROMPTs reads significantly increased in *Dis3*^*cKO*^ oocytes (Fig. 2b-c; Extended Data Fig. 3e-h). By ranking the normalized reads of PROMPTs and genes from RNA-seq, we observed that genes having higher transcription level also have higher PROMPTs level (Fig. 2b). Although the change in abundance of PROMPTs and their associated gene did not correlate in *Dis3*^*cKO*^ mice, there was a tendency for decreased genic transcripts to have increased PROMPTs (Fig. 2d). This suggested a negative effect of PROMPTs on gene level and is consistent with PROMPTs identified in other systems^12,15^.

Although lacking sequence specificity, DIS3 can directly bind and degrade repetitive elements^12^. We hypothesized that DIS3 targeting repetitive elements may underlie pervasive transcripts degradation. We observed that PROMPTs with more SINEs tend to accumulate in greater abundance compared to those with fewer SINEs in *Dis3*^*cKO*^ oocytes (Fig. 2e). Reads in PROMPTs and intergenic regions had stronger SINE association compared to LINE and LTR upon DIS3 depletion (Extended Data Fig. 4a-c), suggesting that DIS3 targeting SINE RNA may coordinate PROMPTs degradation. However, overexpressing synthesized PROMPTs did not cause changes in the nearby genes or PROMPTs, indicating that these free transcripts do not have global cis-or trans-acting effects (Extended Data Fig. 4e-f).

**Figure 2.**
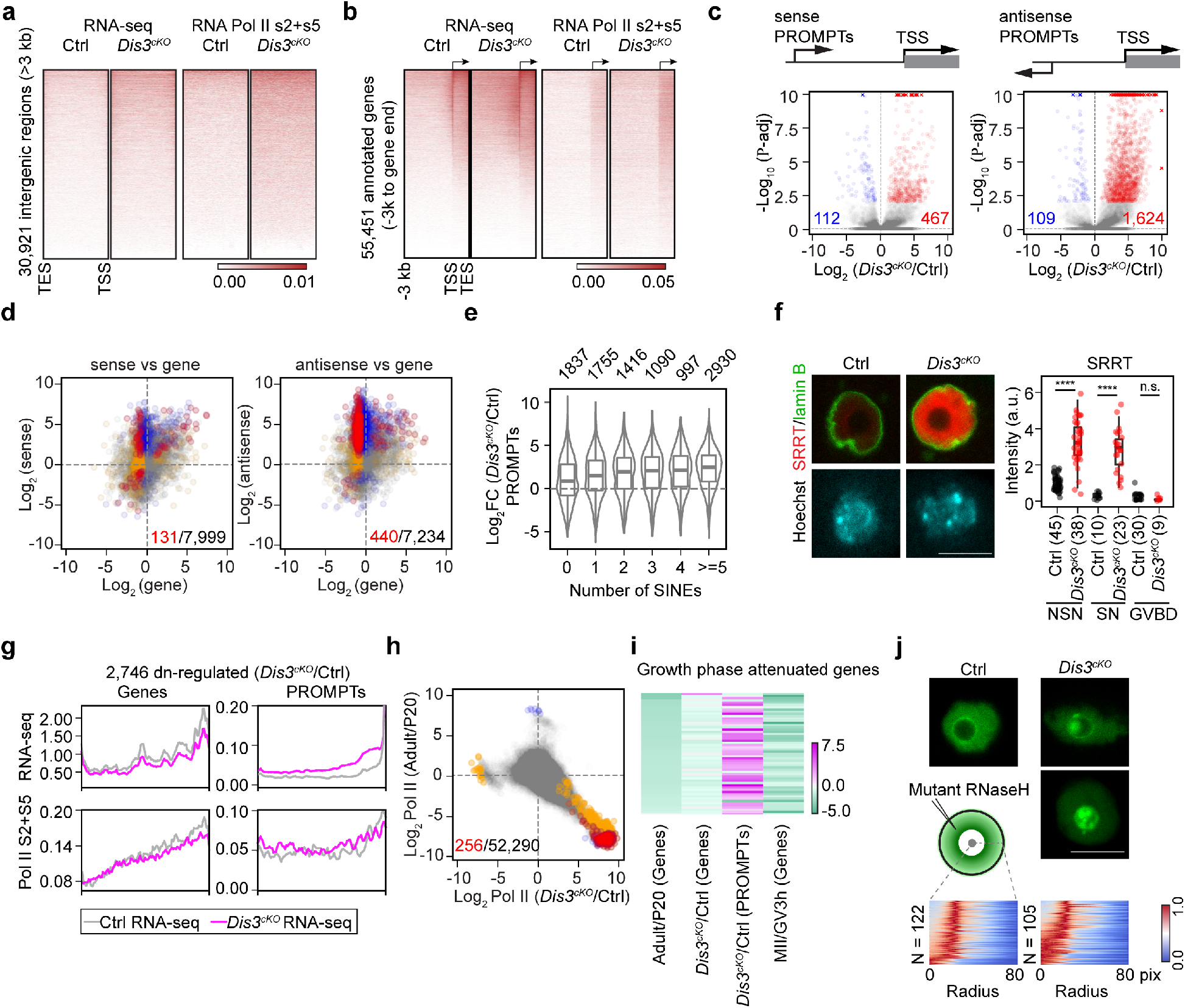
Persistent pervasive transcription and insufficient transcription termination in *Dis3*^*cKO*^ GV oocytes. (**a**) Heatmap of intergenic reads in RNA-seq and in RNA polymerase II Ser2+Ser5 (Pol II S2+S5) CUT&Tag sequencing mapped to 30,812 intergenic regions larger than 3 kb from all annotated genes. All intergenic regions are scaled to the same length. TES and TSS are the end sites and start sites of annotated genes. (**b**) Heatmap of PROMPTs reads (−3 kb to TSS) and scaled genic reads (TSS to end TES) of all 55,451 annotated genes in RNA-seq and in Pol II S2+S5 CUT&Tag sequencing. (**c**) Volcano plots of sense (left) and antisense (right) PROMPTs in *Dis3*^*cKO*^ GV oocytes. PROMPTs of greater or less abundance than Ctrl are labeled in red and blue, respectively (P-adj <0.01). (**d**) Dot plots of changes in sense (left) or antisense (right) PROMPTs abundance vs. changes in downstream genic transcripts at GV stage. A dot represents one gene ID and is color coded if the gene is significantly changed (red: in both, blue: in PROMPTs only, orange: in gene only, gray: not significantly changed). Number of red dots vs. total dots is labeled. (**e**) Violin and boxplot of PROMPTs changing folds grouped by the number of SINEs in their genomic sequence. All antisense PROPMTs that are differentially expressed (independence of significance) are shown. The number above each violin is the number of PROMPTs included in the violin. (**f**) Confocal fluorescence and quantification of SRRT immunostaining at GV stage. **** P = 5.22e-15 (NSN), **** P = 2.64e-10 (SN), n.s., not significant. (**g**) Profiles of read coverage in RNA-seq and Pol II S2+S5 CUT&Tag sequencing of the 2,746 down-regulated genes and their PROMPTs at GV stage. (**h**) Dot plot showing the correlation of Pol II S2+S5 occupancy changes in *Dis3*^*cKO*^/Ctrl at GV stage vs. that in adult/P20 wildtype oocytes. A dot represents one 1000-bp genome bin and is color coded if it is significantly changed (red: in both, blue: in adult/P20 wildtype oocytes only, orange: in *Dis3*^*cKO*^/Ctrl at GV stage only, gray: not significantly changed). (**i**) Heatmap of log_2_ FC values of early attenuated genes (genes that lose transcription from P20 to adult stage) from RNA-seq, including log_2_ FC of genes and PROMPTs in *Dis3*^*cKO*^/Ctrl at GV stage, and log_2_ FC of genes in adult/P20, and MII/GV3h. (**j**) Confocal fluorescence (middle section of the nucleus) of oocytes after microinjecting the cRNA of mutant RNase H (D210N) tagged with V5 and immunostained with V5 antibody. The schematic shows the measurement of radial profile, which is the integrated intensity at each radius step (1.33 pixel). Heatmap shows all the V5 radial profiles in Ctrl and *Dis3*^*cKO*^ oocytes, normalized by min/max values and sorted by the value of the 1^st^ position. Scale bar: 20 μm.

We then tested whether pervasive transcription, rather than free transcripts, has critical impacts on gene transcription. RNA polymerase II Ser2+Ser5 (Pol II Ser2+Ser5) CUT&Tag assays confirmed that increased RNA-seq reads come from the regions occupying more elongating Pol II, indicating that the accumulation of pervasive transcripts not only resulted from insufficient degradation, but also from active transcription (Fig. 2a-b; Supplementary Table 3). The extensive transcription correlates with increased SRRT abundance, a scaffold protein that binds directly to both the RNA exosome complex and newly transcribed RNA to promote transcription termination of short RNAs^16-18^ (Fig. 2f). By profiling the 2,746 down-regulated genes in *Dis3*^*cKO*^ oocytes determined by RNA-seq, we confirmed lower Pol II occupancy in their genic regions but higher Pol II occupancy in their PROMPTs regions (Fig. 2g). A global increase of Pol II occupancy at PROMPTs and intergenic regions vs genic regions also was detected upon DIS3 depletion (Extended Data Fig. 4d).

By further aligning RNA-seq data and Pol II Ser2+Ser5 CUT&Tag sequencing results, we obtained a strikingly negative correlation between genes that normally have decreased transcription during oocyte growth (growth phase attenuated genes) and genes gaining Pol II in *Dis3*^*cKO*^ oocytes at the GV stage. This indicated a persistence of Pol II on chromatin and insufficient transcription termination caused by DIS3 depletion (Fig. 2h). Almost all growth phase attenuated genes had more PROMPTs due to their transcription capability in the genic region, and had reduced gene level due to Pol II relocating to the proximal intergenic region (Fig. 2i). We then microinjected RNase H mutant cRNA to label DNA-RNA hybrid (R-loop), because the pervasive transcription that is not terminated may induce increased R-loops^19,20^. Under normal condition, R-loop signal existed evenly in peri-nucleolar region of the NSN oocytes. However, half of DIS3 depleted oocytes exhibited strong inter-nucleolar or peri-nucleolar R-loop puncta, which resulted in a strong integrated R-loop intensity close to the center of nucleolus (Fig. 2j). Elevated R-loop puncta suggested abnormally persistent R-loop structure caused by insufficient transcription termination^19,21^. In sum, we concluded that gene-proximal intergenic transcripts, such as PROMPTs, generated as byproducts of genic transcription, could support local transcriptional activity and may provide reservoirs for RNA Pol II. Insufficient pervasive transcription termination blocks *Dis3*^*cKO*^ oocytes from resuming meiosis.

Among the down-regulated genes in *Dis3*^*cKO*^ oocytes, we observed those that encode histone modifiers (Supplementary Table 4) and regulate GV oocytes chromatin remodeling^2,22-24^. Immunostaining of H3K4me3 and H3K9me3 indicated unchanged levels in *Dis3*^*cKO*^ oocytes (Extended Data Fig. 5a-d). However, H3K27me3 was significantly increased in both NSN and SN *Dis3*^*cKO*^ oocytes (Fig. 3a), probably due to the decrease of demethylases KDM6A/B (Fig. 3b; Supplementary Table 4). Gain of H3K27me3 may underlie the significant decrease of LTR transcripts in *Dis3*^*cKO*^ oocytes (Extended Data Fig. 4a-b), because H3K27me3 specifically represses retrotransposons^25,26^. In addition, increased H3K27me3 could be partially rescued by overexpressing *Kdm6a/b* (Fig. 3c). CUT&Tag assays suggested an overall increase of H3K27me3 peak numbers in *Dis3*^*cKO*^ oocytes in both genic and intergenic regions (Fig. 3e; Supplementary Table 5). Next, we examined the co-regulation of H3K27me3 and Pol II. Specifically, the 4,584 genes up-regulated from GV to GV3h in *Dis3*^*cKO*^ oocytes had higher Pol II and lower genic H3K27me3 occupancy, consistent with their continuing production of transcripts. We then aligned our H3K27me3 peaks with known polycomb-associating domains (PADs, having high H3K27me3) and inter-PAD (iPAD, having lower H3K27me3) regions defined by local Hi-C analyses of mouse oocytes^6^. Interestingly, change of H3K27me3 in *Dis3*^*cKO*^ oocytes happened as predicted: 3/4 of increased peaks localize in PAD region, while 2/3 of decreased peaks localize in iPAD region with very few peaks crossing PAD/iPAD boundaries (Fig. 3f-g; Extended Data Fig. 5f). This indicated that gain of H3K27m3 in *Dis3*^*cKO*^ oocytes happened in a pre-defined manner. In addition, we observed almost no overlap between significantly changed H3K27me3 and Pol II Ser2+Ser5 1000-bp genome bins, suggesting the regulation of the active transcription and repressive chromatins do not locally interfere with each other, but globally occupy complementary chromatin domains in oocyte development (Fig. 3f-h).

**Figure 3.**
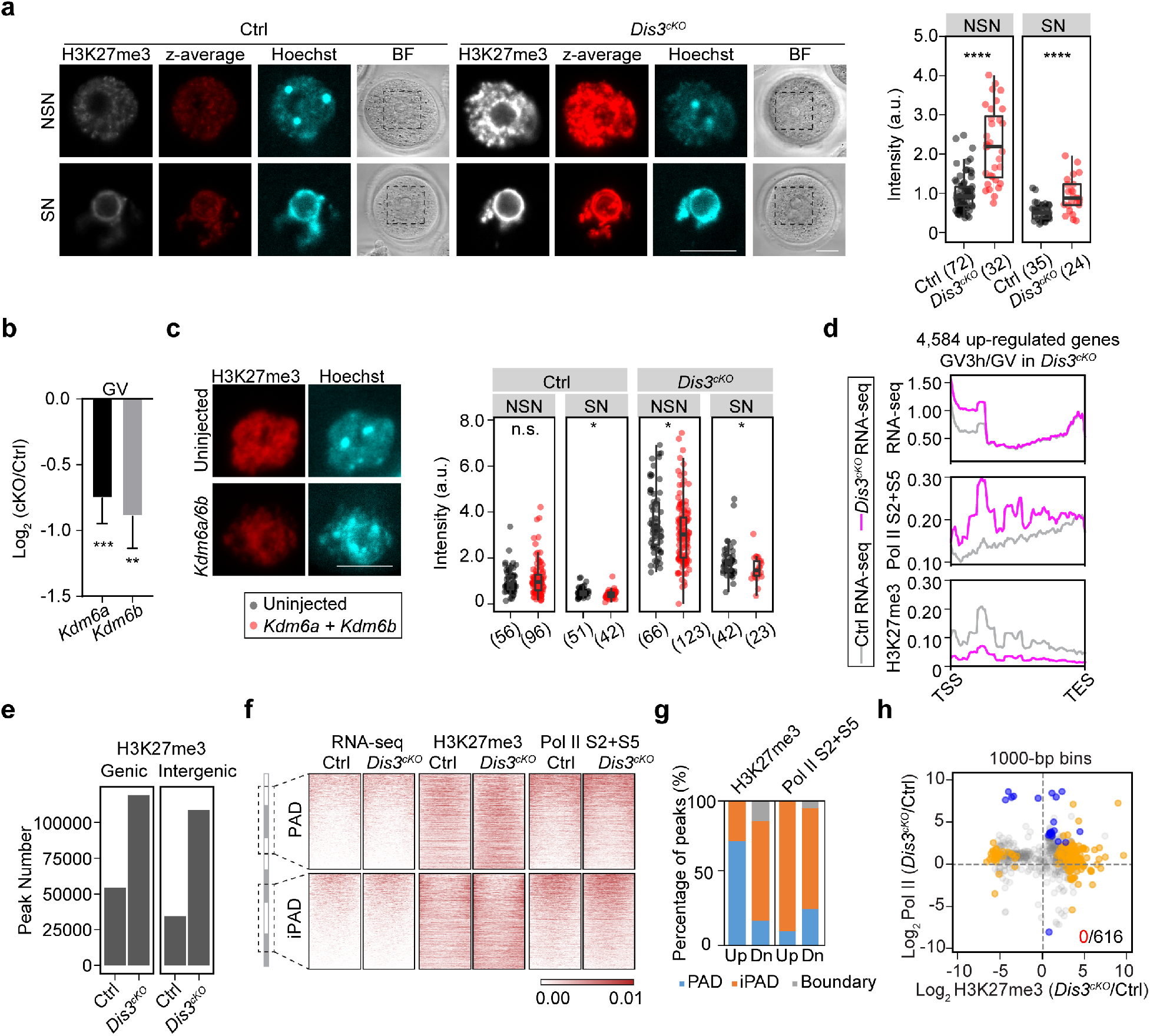
*Dis3*^*cKO*^ oocytes have excessive H3K27me3 due to decreased *Kdm6a/b*. (**a**) Confocal fluorescence and quantification of H3K27me3 immunostaining of NSN and SN oocytes at GV stage. **** P = 3.14e-08 (NSN), **** P = 1.21e-04 (SN). Z-average, average intensity projection. (**b**) *Kdm6a* and *Kdm6b* transcript level decreases in *Dis3*^*cKO*^ oocytes. *** P*-*adj < 0.001, ** P*-*adj < 0.01. (**c**) Confocal fluorescence and quantification of H3K27me3 immunostaining after microinjection of *Kdm6a/b* cRNAs. * P = 0.026 (Ctrl-SN), * P = 0.018 (*Dis3*^*cKO*^-NSN), * P = 0.031 (*Dis3*^*cKO*^-SN). (**d**) Profiles of read coverage from RNA-seq and CUT&Tag sequencing of Pol II S2+S5 and H3K27me3 of the 4,586 up-regulated genes in *Dis3*^*cKO*^ from GV to GV3h stage. (**e**) H3K27me3 peak distribution in Ctrl and *Dis3*^*cKO*^ oocytes at GV stage in genic and intergenic regions. (**f**) Heatmap of reads (from RNA-seq) and peaks (from CUT&Tag sequencing of H3K27me3 and Pol II S2+S5) localized in PAD (polycomb associated domain) and iPAD (inter-PAD) defined by local Hi-C analyses of mouse oocytes^6^. (**g**) Percentage of differential peaks in PAD, iPAD and boundaries. (**h**) Correlation plot of H3K27me3 and Pol II S2+S5 occupancy changes in *Dis3*^*cKO*^/Ctrl at GV stage. A dot represents one 1000-bp genome bin and is color coded (red: in both, blue: in Pol II S2+S5 only, orange: in H3K27me3 only, gray: not significantly changed). Number of red dots vs. total dots is labeled. Scale bar: 20 μm.

We previously reported that depletion of EXOSC10 results in growth-to-maturation transition defects^3^. Given that both proteins participate in the RNA exosome complex, we crossed the two lines to establish double oocyte-specific knockout mice (dcKO). As expected, dcKO females have greatly reduced fertility (Fig. 4a). As early as the mid-growth stage (P20), a reduction of oocyte size was observed in the dcKO, while oocytes at the same stage in *Exosc10*^*cKO*^ and *Dis3*^*cKO*^ ovaries appeared normal (Fig. 4b; Extended Data Fig. 6a-b). Consequently, the ovary size became significantly smaller from ∼6 weeks in the dcKO females (Extended Data Fig. 6c-e). Clustering of RNA-seq samples at P20 and adult GV stage showed that the transcriptome of P20 dcKO oocytes were distinct from wildtype and single mutant oocytes (Fig. 4c; Supplementary Table 6). dcKO oocytes also had retained excessive intergenic transcripts and PROMPTs (Fig. 4d-f; Extended Data Fig. 6f-g).

**Figure 4.**
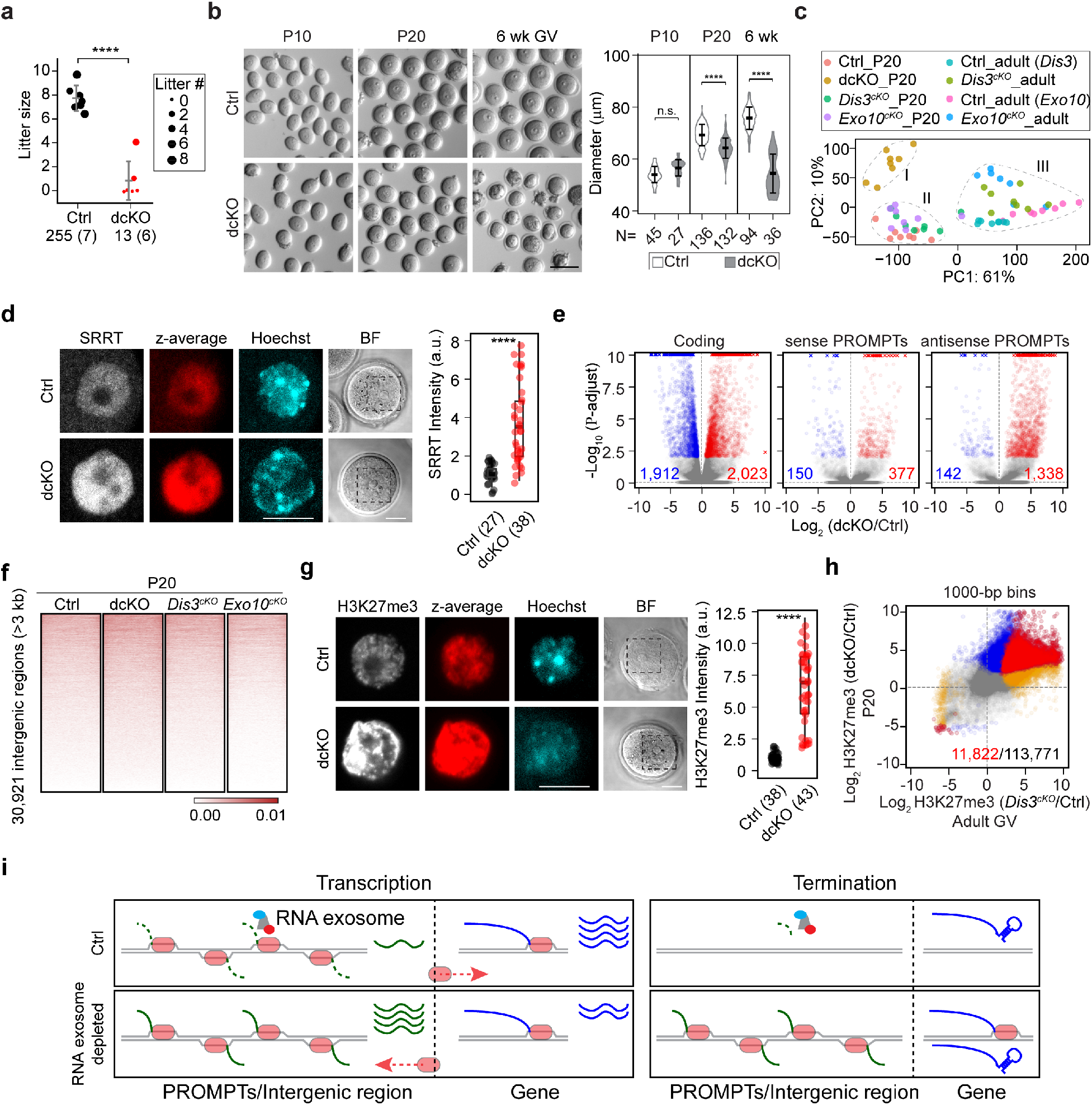
EXOSC10 functions synergistically with DIS3 in oocyte development and in pervasive transcript degradation. (**a**) Dot plot of fertility record. Each dot represents one female from a harem breeding of Ctrl and dcKO females with wildtype males. Dot sizes represent the average litter number. The number of pups and females of each genotype is indicated below each group. The horizontal lines represent the mean and standard deviation. **** P *=* 1.66e-06. (**b**) Oocytes dissected from follicles in Ctrl and dcKO females at postnatal 10 days (P10), 20 days (P20) and 6 weeks (6 wk). Oocyte diameter is shown on right. The horizontal lines represent the mean and standard deviation. **** P *=* 7.16E-22 (P20), **** P *=* 2.65E-20 (6 wk). Scale bar: 100 μm. The number of oocytes included is shown below each group. (**c**) Dot plot showing PCA of all RNA-seq samples. I, II, III were three clusters obtained by K-means clustering (define k=3) based on the PCA. (**d, g**) Confocal fluorescence and quantification of SRRT (d) and H3K27me3 (g) of P20 oocytes. Scale bar: 20 μm. **** P *=* 6.15e-10 (d), **** P *=* 2.00e-15 (g). The number of oocytes included is shown below each group. (**e**) Volcano plots of genic transcripts (left), sense PROMPTs (middle) and antisense PROMPTs (right) changes in dcKO/Ctrl. (**f**) Heatmap of intergenic reads in RNA-seq in Ctrl, dcKO, *Dis3*^*cKO*^ and *Exosc10*^*cKO*^ oocytes at P20. (**h**) Correlation of H3K27me3 changes in *Dis3*^*cKO*^/Ctrl of adult GV oocytes and in dcKO/Ctrl of P20 oocytes. A dot represents one 1000-bp genome bin and is color coded if it is significantly changed (red: in both, blue: in dcKO/Ctrl of P20 oocytes only, orange: in *Dis3*^*cKO*^/Ctrl of adult GV oocytes only, gray: not significantly changed). The number of red dots vs. total dots is labeled. (**i**) Schematic of the RNA exosome function in clearing pervasive transcription and transcripts to serve as a reservoir of RNA polymerase. In the absence of RNA exosome, persistent pervasive transcription sequesters RNA polymerase to reduce gene transcription while simultaneously maintaining local transcription activity.

We observed a high similarity in gain of H3K27me3 occupancy in *Dis3*^*cKO*^ GV oocytes and P20 dcKO oocytes (Fig. 4g-h), suggesting a synergistic effect of DIS3 and EXOSC10 in protecting the transcriptome integrity in oocyte development. First, the two RNases do not have genetic complementarity on the transcriptional level (Extended Data Fig. 7a). Secondly, though DIS3 depletion resulted in much more PROMPTs accumulation, degradation of more than half of the PROMPTs depended on both RNases and accumulation of these PROMPTs is observed despite DIS3 or EXOSC10 depletion (Extended Data Fig. 7b-c). Interestingly, the positive correlation between DIS3 and EXOSC10 happens more significantly in PROMPTs change rather than genes change (Extended Data Fig. 7b). Thirdly, the short PROMPTs RNA had similar accumulation after DIS3 or EXOSC10 depletion (Extended Data Fig. 7d-e).

In summary, our study suggests that the RNA exosome protects genome stability to allow meiosis resumption by degradation-coupled transcription termination during late oocyte growth. The otherwise undetectable pervasive intergenic transcription serves as RNA Pol II reservoirs and share similar transcription activity with the gene. Upon RNA exosome depletion, intergenic pervasive transcripts are increased due to both persistent transcription and insufficient degradation, which could subtract Pol II from genic regions while maintaining local active state of transcription (Fig. 4i). We highlight the function of RNA exosome in a unique cellular context of global transcription termination and chromatin compaction in fully-grown oocytes. Our study incorporates the role of histone demethylases in actively maintaining the H3K27me3 pattern and chromatin compartmentalization of active or repressive transcription in mouse oocytes.

## Methods

### Mice generation

Mice were maintained in compliance with the guidelines of the Animal Care and Use Committee of the National Institutes of Health (Animal study protocol K018LCDB21).

The *Dis3* floxed allele was generated by CRISPR/Cas9. Two loxP sequences were inserted surrounding exons 3-5 of *Dis3* genomic region. A BamHI restriction site is designed next to the left loxP and an EcoRI is next to the right loxP to facilitate genotyping. Each guide RNA (gRNA) was synthesized by *in vitro* transcription using a MEGAshortscript™ T7 Transcription Kit (ThermoFisher AM1354) and purified by a MEGAclear™ Transcription Clean-Up Kit (ThermoFisher AM1908). The templates for gRNA *in vitro* transcription were synthesized as single stranded DNA (ssDNA) by Integrated DNA Technologies (IDT). The two homology-directed repair templates (HRTPs) were synthesized as ssDNA by IDT. *Cas9* mRNA was synthesized by *in vitro* transcription using mMESSAGE mMACHINE™ T7 ULTRA Transcription Kit (ThermoFisher AM1345) from plasmid MLM3613, which was a gift from Keith Joung (Addgene plasmid # 42251)^27^ and purified by the MEGAclear™ Transcription Clean-Up Kit. All DNA, RNA, genotyping primer sequences are available in Supplementary Table 7.

To prepare mouse 1-cell embryos for microinjection, hormonally stimulated B6D2_F1_ female mice were mated to B6D2_F1_ stud males. 1-cell zygotes were flushed from oviducts into M2 medium (CytoSpring M2114), treated with 0.5 mg/ml Hyaluronidase to remove cumulus cells, cultured to the pronuclear stage and microinjected with the mixed injection solution, including the two gRNAs (50 ng/μl each), two HRTPs (100 ng/μl each) and *Cas9* cRNA (100 ng/μl). After *ex vivo* culture in KSOM medium (CytoSpring K0113) for 1 day, the injected 2-cell embryos were transferred into the oviducts of 0.5-day post coitus pseudopregnant ICR females. The obtained *Dis3* floxed mice were crossed to *Zp3-cre*^*28*^ mice to obtain oocyte-specific conditional *Dis3* knockout mice (*Dis3*^*cKO*^). *Exosc10;Dis3* double conditional knockout mice (dcKO) were generated by crossing *Exosc10* flox/flox mice, *Dis3* flox/flox mice and *Zp3-cre* mice.

### Mouse oocyte collection, culture, and microinjection

Ovaries were dissected from female mice (2-4 months old or P20) into M2 medium containing milrinone (2.5 μM) and pierced mechanically by 30-gauge needles. Fully-grown oocytes were washed with milrinone-M2 medium 10 times to remove granulosa cells. For *ex vivo* oocyte maturation, GV oocytes were washed with M2 medium without milrinone 10 times and cultured in M2 medium at 37 °C with 5% CO_2_ for 3 hours (to GVBD) or for 14-16 hours (to mature eggs). During microinjection, oocytes were held in milrinone-M2 medium. The microinjection system is composed of an Eppendorf Femtojet 4i Injector, an inverted microscope (Zeiss) and a pair of Eppendorf Transferman NK2 micro-manipulators. Microinjection needles were prepared from Sutter Instrument Borosilicate glass with filament (BF150-75-10), using a Sutter Flaming/Brown Micropipette Puller. An Eppendorf Microloader (930001007) was used to load the injection solution into a needle, and an Eppendorf VacuTip I (5195000036) was used to hold and transfer oocytes during injection. The injector was set in Auto mode, with the injection time (ti) 0.1 s, the compensation pressure (Pc) 15, and the injection pressure (Pi) 200.

### cRNA synthesis for microinjection

cRNAs were synthesized by mMESSAGE mMACHINE™ SP6 Transcription Kit (ThermoFisher AM1340) or mMESSAGE mMACHINE™ T7 ULTRA Transcription Kit (ThermoFisher AM1345). The templates were linearized from plasmids or PCR amplified cDNA. The synthesized RNAs were purified with the MEGAclear™ Transcription Clean-Up Kit and diluted for microinjection. The templates for *Kdm6a/Kdm6b* cRNA synthesis were from pCS2-UTX-F and pCS2-Jmjd3-F, which were gifts from Dr. Kai Ge (Addgene plasmid # 17438, and # 17440)^29^. The template for *RNaseH D210N* was from ppyCAG-RNaseH1-D210N, which was a gift from Dr. Xiang-Dong Fu (Addgene plasmid # 111904)^30^. The synthesized PROMPTs sequences are in Supplementary Table 7. Each PROMPT sequence was surrounded by a T7 and a T3 sequence at the 5’ and 3’ end, respectively, for template amplification. The injected PROMPTs transcripts were synthesized using mMESSAGE mMACHINE™ T7 ULTRA Transcription Kit and purified with the MEGAclear™ Transcription Clean-Up Kit.

### Immunofluorescence and confocal microscopy

Oocytes were fixed in 2% paraformaldehyde (ThermoFisher 50-980-487) diluted in PBS containing 0.1% Tween-20 (Sigma P1379-25ML) for 30 min at 37 °C. Fixed oocytes were washed with PBVT (PBS, 3 mg/ml polyvinylpyrrolidone-40 and 0.1% Tween-20) once and permeabilized with 0.5% Triton X-100 in PBS for 30 minutes at room temperature. Oocytes were washed with PBVT twice and blocked with 5% normal goat serum (Sigma-Aldrich G9023-10ML) diluted in PBVT for 1 hour at room temperature and then incubated with primary antibody overnight at 4 °C. On the second day, oocytes were washed with PBVT (4X, 20 minutes) and incubated with fluorescence conjugated secondary antibody overnight at 4 °C. On the third day, oocytes were washed with PBVT (4X, 20 minutes), stained with Hoechst (1:5000 diluted in PBS) for 15 minutes at room temperature and immediately transferred into PBS for confocal microscopy (Carl Zeiss LSM 780, ZEN SP5). During imaging, oocytes were held in µ-Slide 18-Well Glass Bottom chambered glass coverslip (Ibidi 81817). Images were processed using Image J (Fiji).

### Antibodies and staining reagents

DIS3 (1:200, Abcam 223767); GM130 (1:200, BD Transduction Laboratories 610823); lamin B1 (B-10) (1:200, Santa Cruz sc-374015); lamin A/C (4C11) (1:200, CST 4777T); α-tubulin (1:200, Sigma T5168); phospholamin A/C (Ser22) (1:200, CST 2026); RAB5 [EPR21801] (1:500, Abcam ab218624); phosphoCDK1 (Thr14, Tyr15) (17H29L7) (1:200, ThermoFisher 701808); RAB7 (1:500, Abcam ab137029); H3K4me3 (1:500, CST mAb 9751); H3K9me3 (1:500, CST mAb 13969); H3K27me3 (1:500, CST mAb 9733); SRRT (1:200, Aviva Systems Biology Corporation OAGA06370); RNA polymerase II CTD Panel (RNA pol II CTD, phospho S2, phospho S5) (ab103968); V5 Tag Monoclonal Antibody DyLight 488 conjugated (1:200, Invitrogen MA5-15253-D488); Flag antibody (ThermoFisher F1804); Alexa Fluor™ 555 Tyramide SuperBoost™ Kit, goat anti-rabbit IgG (ThermoFisher B40923); goat anti-rabbit IgG (H+L) cross-adsorbed secondary antibody, Alexa Fluor 546 (1:500, Invitrogen A-11010); goat anti-mouse IgG (H+L) cross-adsorbed secondary antibody, Alexa Fluor 647 (1:500, Invitrogen A-21235); DAPI (Sigma D9542-1MG).

### Ovary histology

Ovaries were dissected from female mice at desired stages into PBS and surrounding tissues were cleared. Then ovaries were fixed in newly prepared 2.5% glutaraldehyde (VWR 100504-790) and 2.5% paraformaldehyde (Fisher Scientific 50-980-492) in 0.083 M sodium cacodylate buffer (pH 7.2) (VWR 102090-966) at 4 °C for 3-5 hrs. Ovaries were washed with 0.1 M sodium cacodylate buffer (pH 7.2) and stored at 4 °C overnight. Then ovaries were transferred into 70% ethanol and stored at 4 °C. 2 μm plastic sectioning and periodic acid– Schiff staining were performed by American HistoLab. The imaging of sections was performed in tiles and stitched together with Image J (Fiji).

### Translation activity detected by Click-iT HPG assay

Translation activity in oocytes was detected by the Click-iT® HPG Alexa Fluor® 594 Protein Synthesis Assay Kit (ThermoFisher C10429). Oocytes were cultured in M2 medium containing milrinone (2.5 μM) and HPG (50 μM) at 37 °C for 30 min. Then oocytes were washed twice with PBS, fixed with 2% paraformaldehyde, washed twice with 3% BSA-PBS and detected with a Click-iT HPG kit per the manufacture’s protocol. Oocytes were imaged by confocal microscopy and intensity was measured with Image J (Fiji).

### Single oocyte RNA-seq

Poly(A)-based and RiboMinus RNA-seq of single oocytes were performed as described^3,31^.

Briefly, for poly(A)-based RNA-seq, single oocytes at the desired stages were collected individually in 2.5 μl RLT Plus (Qiagen 1053393). Each oocyte lysis was supplemented with 1 μl of the 1:10^5^ diluted ERCC spike-in mix (ThermoFisher 4456740). Poly(A) RNA was isolated by oligo (dT) beads, reverse transcribed, amplified for 16 cycles, purified with AMPure beads (AMPure XP, Beckman Coulter, A63880), and evaluated by Bioanalyzer 2100 (Agilent). Quantified cDNAs were used to construct sequencing libraries using the Nextera DNA Sample Preparation Kits (Illumina).

RiboMinus RNA-seq libraries were prepared with the Ovation® SoLo RNA-Seq Library Preparation Kit (Tecan # 0501-32). Each oocyte was collected in lysis buffer, supplemented with 1 μl of the 1:10^5^ diluted ERCC spike-in mix and 1 μl (50 μM) of random hexamer primer (ThermoFisher 3005) for first strand cDNA synthesis. cDNAs were processed and amplified for 16 cycles, purified, and analyzed by Bioanalyzer 2100 (Agilent). The ribosomal DNA was depleted, and the remaining DNA was amplified, purified, pooled and analyzed by Bioanalyzer 2100. The sequencing was performed by the NIDDK Genomic Core Facility using the HiSeq 3000 Sequencing System (Illumina).

### RNA-seq analysis

The RNA-seq FASTQ files were aligned with STAR (v2.7.8a)^32^ to obtain BAM files. HTSeq (v0.11.4)^33^ was used for calculating the number of reads mapped to genes, PROMPTs or intergenic regions using the corresponding reference files. The distribution of reads of UTR, intronic, intergenic and coding regions were calculated by Picard Tools (v2.26.9). The differential analysis was performed by DESeq2 (v1.24.0)^34^ in R and ERCC molecules were used for library size normalization across samples. Key scripts used to analyze the RNA-seq data are available from GitHub: https://github.com/Di-aswater/public-scripts-Dis3cKO

### CUT&Tag experiment

CUT&Tag was performed as described^35^. Briefly, fresh denuded oocytes at the desired stages (adult or P20) were collected and washed with M2 medium containing milrinone (2.5 μM) 10 times to clear the surrounded somatic cells. Each treatment (antibody and stage) had three biological replicates and each replicate had 80-150 oocytes. Oocytes were washed twice with EmbryoMax® Acidic Tyrode’s Solution (EMD Millipore MR-004-D) to remove zonae pellucidae and washed twice with M2 medium containing milrinone. Oocytes were washed once with PBS, transferred into 100 µL wash buffer prepared as described^35^ at room temperature and incubated with pre-treated concanavalin A beads. Primary antibodies used include H3K27me3 (Cell Signaling Technology 9733) and RNA polymerase II Ser2+Ser5 (Cell Signaling Technology, Rpb1 CTD Antibody Sampler Kit 54020). All primary antibodies were diluted 1:100 in antibody buffer and rabbit IgG was used as the negative control. After incubation at 4 °C overnight, oocytes were incubated with a secondary antibody (guinea pig anti-rabbit ABIN101961) for 2 hrs at room temperature. Oocytes were washed twice with Dig-wash buffer and incubated with pA-Tn5 (Epicypher) for 1 hr at room temperature. Oocytes were washed twice with Dig-300 buffer and incubated in tagmentation buffer at 37 ºC for 1 hr on a thermomixer (500 rpm). Tagmentation was stopped by adding 4 µL 0.5M EDTA, 1 µL 10% SDS and 1 µL 20 mg/mL Proteinase K to each sample (per 100 µL tagmentation solution). The reaction solution was mixed and incubated for 1 hr at 55 ºC on a thermomixer. ConA beads were removed, and the supernatant was transferred into a new tube and incubated at 70 °C for 20 min to inactivate Proteinase K. DNA was extracted using AMPure beads (AMPure XP, Beckman Coulter, A63880). All purified DNA was barcoded individually for 17 cycles of PCR amplification. Replicates of the same antibody and stage were mixed for purification using AMPure beads. All samples were pooled for paired-end sequencing by the NIDDK Genomic Core Facility using the NextSeq High output mode.

### CUT&Tag analysis

Each CUT&Tag original fastq file was randomly sampled 3 million reads by Seqtk tools (v1.3) for subsequent downstream analyses^36^. Reads were processed without deduplication. Peaks were called using SEACR (v1.2)^37^ and the IgG samples were implemented as the background control. Peak number for each sample was determined from the overlap peaks of at least two replicates. In differential analysis, because of the obvious overall increase of H3K27me3 indicated by immunofluorescence assays, the ChIPseqSpikeInFree^38^ package was employed for size factor evaluation and normalization between samples. The obtained size factors were used in DESeq2 (v1.24.0) for differential analysis of consensus 1000-bp genome bins. The differential peaks were further computed against genes, intergenic regions, and PROMPTs reference files to get the read coverage on each region using BEDTools (v2.30.0). Heatmaps and profiles were produced using deepTools (v3.5.1)^39^.

### Intergenic region analysis

Intergenic regions were extracted from GRCm38 using BEDTools (v2.30.0). Only the intergenic regions larger than 3 kb were used for subsequent analyses. The PROMPTs reference file was generated by extracting the -3 kb to -1 bp upstream of each gene and excluding the ones that were overlapped with other annotated features. BAM files were used for counting against the intergenic region or PROMPTs reference file to get the intergenic reads or PROMPTs reads for downstream differential analysis.

To plot heatmaps, BAM files of the same genotype/stage were merged. BAM coverage files normalized by CPM were generated from the merged BAM files using deepTools (v3.5.1). The BAM coverage files were then processed by deepTools against the corresponding reference files and were further used to plot heatmaps and profiles.

### Repetitive element analysis

The *repeat_masker_mm10*.*fa*.*out* reference file was downloaded from *repeatmasker*.*org*, from which the reference files of interspersed repeats including SINE, LINE, LTR were generated using BEDTools (v2.30.0). The SINE/LINE/LTR reference files were intersected with PROMPTs and the intergenic reference files to obtain the PROMPTs-located or intergenic SINE/LINE/LTR and to calculate the number of each element on each PROMPTs or intergenic region.

### Quantitative RT-PCR

20-30 oocytes were collected for RNA extraction using RNeasy Plus Micro Kit (Qiagen 74034). Purified RNA was used for reverse transcription using SuperScript™ II Reverse Transcriptase kit (ThermoFisher 18064014) and random hexamer (ThermoFisher 48190011). qRT-PCR was performed by iTaq Universal SYBR Green Supermix (Bio-Rad #1725121) and QuantStudio 6 Flex Real-Time PCR System (ThermoFisher). Each quantitative RT-PCR assay was repeated 2-3 times, with each time having at least 3 biological replicates and each biological replicate having 3 technical replicates. The error bars represent standard deviation of the 3 technical replicates of each sample. The bar graphs were from one representative assay. The primers used for qPCR are in Supplemental Table 7.

### Statistics and reproducibility

In all statistical comparisons involved in this study, including fertility, immunofluorescence intensities, ovary weights, P values were calculated using two-tailed Student’s *t*-test when two samples were compared, and were calculated using one-way ANOVA when more than two samples were compared.

## Data availability

The sequencing data reported in this study has been deposited in the Gene Expression Omnibus website with accession code GSE196331. All reference files and scripts are available at https://github.com/Di-aswater/public-scripts-Dis3cKO

## Additional information

**Supplementary Table 1**. Differentially expressed gene lists and PROMPTs lists obtained from poly(A) single-oocyte RNA-seq at GV, GV3h and MII stages. The table contains 14 sheets, including 7 gene lists and 7 PROMPTs lists of GV_cKO vs. GV_ctr, GV3h_cKO vs. GV3h_ctr, MII_cKO vs. MII_ctr, GV3h_ctr vs. GV_ctr, MII_ctr vs. GV3h_ctr, GV3h_cKO vs. GV_cKO and MII_cKO vs. GV3h_cKO.

**Supplementary Table 2**. Differentially expressed gene lists and PROMPTs lists obtained from RiboMinus single GV oocyte RNA-seq. The table contains 3 sheets, include 1 of genes, 1 of sense PROMPTs and 1 of antisense PROMPTs of GV_cKO vs. GV_ctr.

**Supplementary Table 3**. CUT&Tag of RNA polymerase II Ser2 + Ser5 (Pol II S2+S5) differentially occupied 1000-bp genome bins. The table contains 2 sheets, including adult_GV_*Dis3*^*cKO*^ vs. adult_GV_ctr and adult_GV_ctr vs. P20_ctr. All 1000-bp genome bins examined are shown.

**Supplementary Table 4**. Summary of histone modification genes obtained from RiboMinus RNA-seq differential analysis.

**Supplementary Table 5**. CUT&Tag of H3K27me3 differentially occupied 1000-bp genome bins. The table contains 2 sheets, including adult_GV_*Dis3*^*cKO*^ vs. adult_GV_ctr and P20_dcKO vs. P20_ctr.

**Supplementary Table 6**. Differentially expressed gene lists and PROMPTs lists obtained from poly(A) single-oocyte RNA-seq at P20 stage. The table contains 9 sheets, including 3 gene lists, 3 sense PROMPTs lists and 3 antisense PROMPTs lists of dcKO_P20 vs. ctr_P20, *Dis3*^*cKO*^_P20 vs. ctr_P20 and *Exosc10*^*cKO*^_P20 vs. ctr_P20.

**Supplementary Table 7**. All oligos, primers and synthesized genes used in this study.

