## Supplementary figures for "Persistent pervasive transcription in RNA exosome depleted oocytes results in loss of female fertility"

#### Supplementary Materials

#### Extended Data Fig. 1

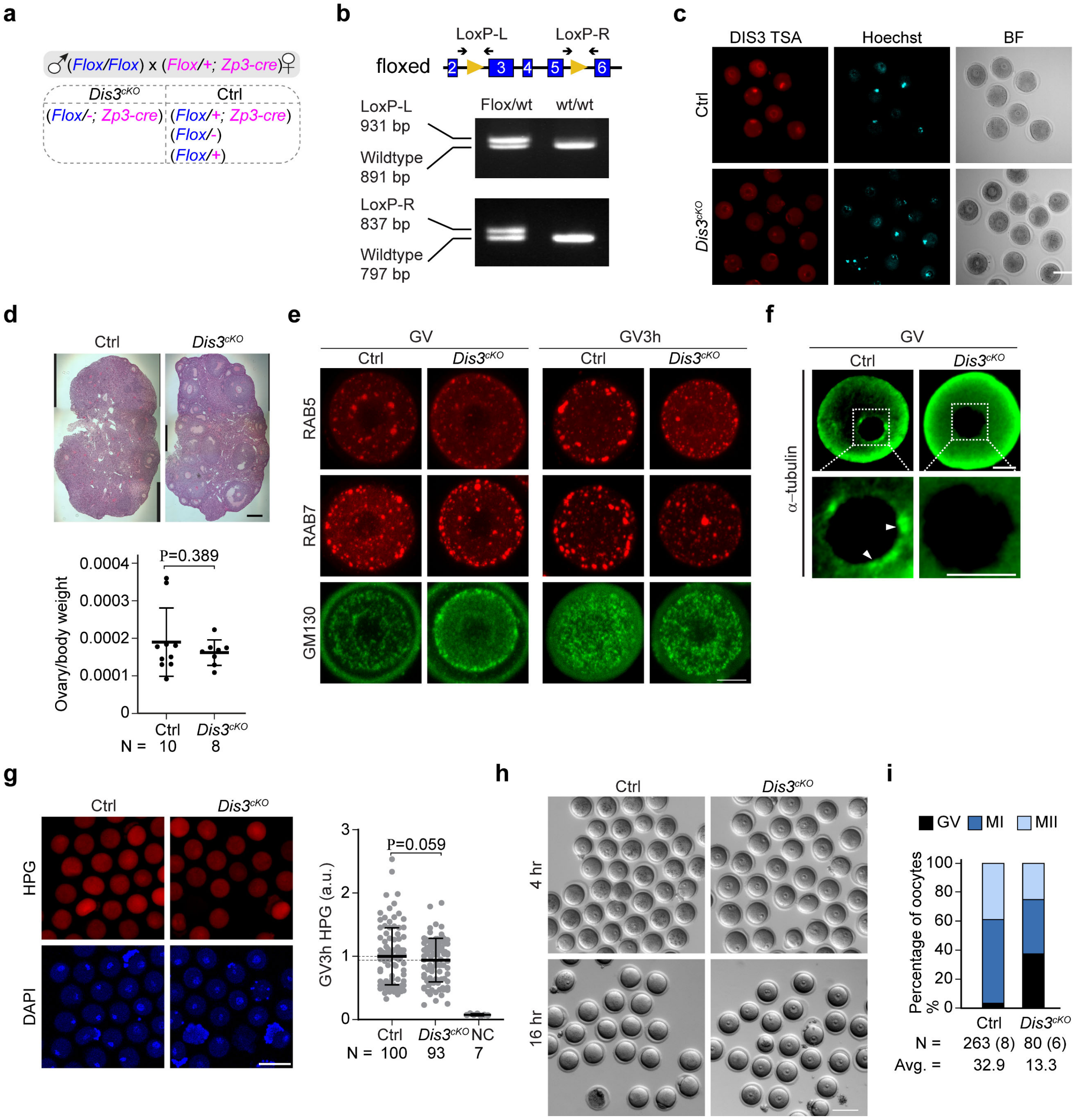


**Extended Figure 1. *Dis3*$^{cKO}$ oocytes are arrested at the GV stage.** (**a**) Mating strategy to obtain oocyte-specific knockouts of *Dis3*. The paternal allele is labeled in blue, and the maternal allele is labeled in magenta. Note that the maternal *Dis3* floxed allele will become a deletion allele (-) during oocyte growth due to the expression of *Zp3-cre* in *Dis3*$^{cKO}$ growing oocytes. (**b**) Schematic of genotyping strategy of two loxPs and an example of genotyping results from a mouse having a floxed allele. loxP-L: the left loxP site; loxP-R: the right loxP site. For both left and right loxP sites, the floxed allele is larger than wildtype allele due to the insertion of a loxP. (**c**) DIS3 antibody staining followed by Tyramide Signal Amplification (TSA) of Ctrl and *Dis3*$^{cKO}$ GV oocytes. (**d**)

Ovary histology (top) and normalized ovary weight (bottom) of Ctrl and *Dis3*$^{cKO}$ females at 12 weeks. (**e**) Confocal fluorescence of RAB5, RAB7 and GM130 immunostaining at GV and GV3h stage. (**f**) Confocal fluorescence of $\alpha$-tubulin immunostaining at GV stage. White triangles indicate the peri-nuclear puncta of $\alpha$-tubulin. (**g**) Overall translation activity detected by HPG labeling at GV and GV3h stages. NC: negative control that were not incubated with HPG. The number of oocytes tested is labeled below each group. (**h**) Live imaging of oocytes at 4 hours and 16 hours after *ex vivo* culture of GV oocytes. (**i**) Percentage of GV, meiosis I (MI) and meiosis II (MII) oocytes obtained from ovulation of Ctrl and *Dis3*$^{cKO}$ females at 12 weeks. The number of oocytes and females is labeled below each group. Avg., average number of ovulated oocytes per female mouse. Scale bar: 20 $\mu$m in e, f, 100 $\mu$m in c, d, g, h.

### Extended Data Fig. 2

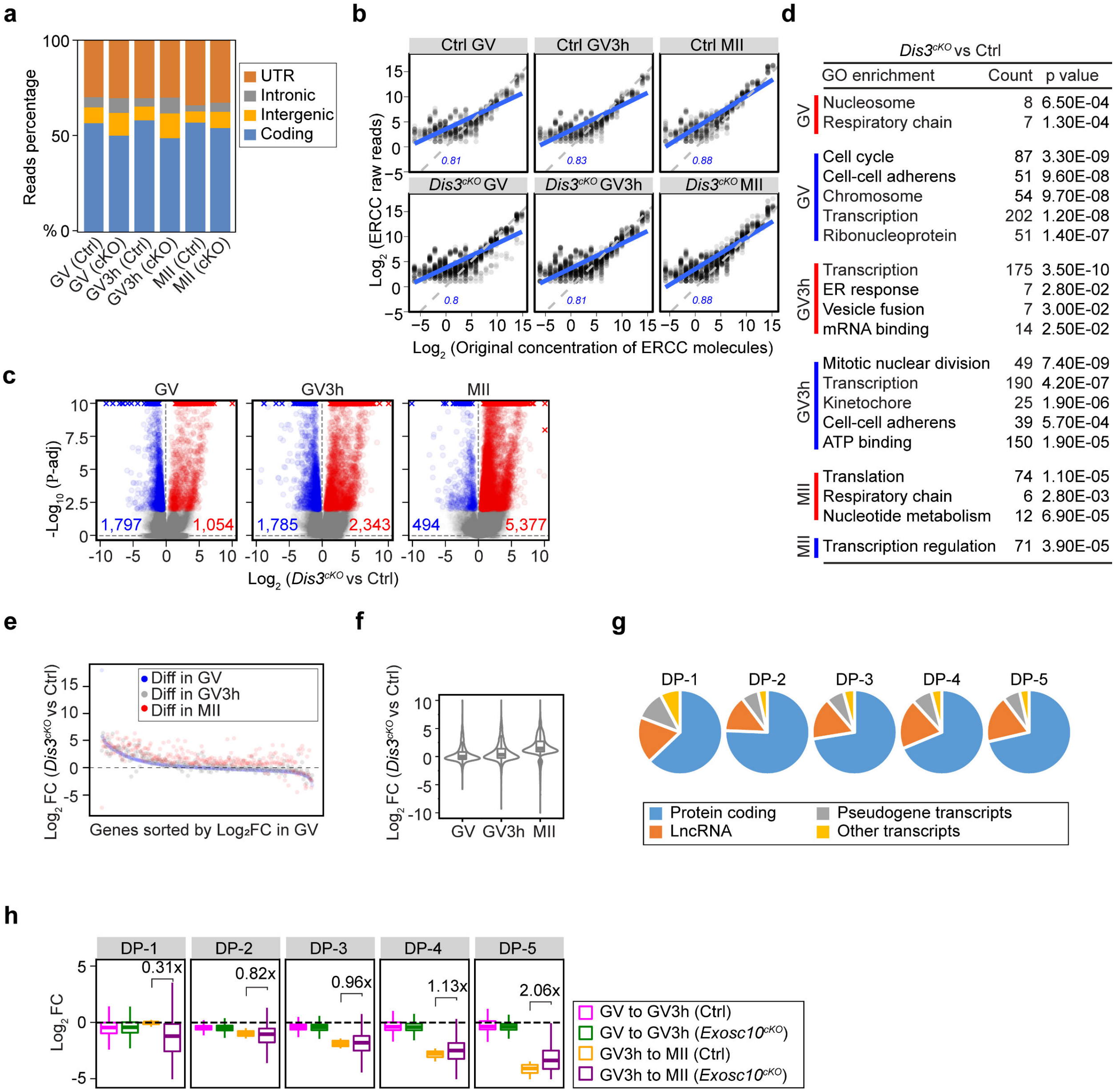


**Extended Figure 2. Transcriptome is disrupted in *Dis3$^{cKO}$* oocytes during meiotic maturation.** (**a**) Percentage of reads mapped to UTR, intergenic, intronic and coding regions in all samples. Samples of the same condition were merged. (**b**) Correlation plots of the original concentration and gene count of the ERCC spike-in molecules in RNA-seq results. Blue lines and blue numbers: linear regression and coefficient of correlation. (**c**) Volcano plots of differentially expressed genes at GV, GV3h and MII stages. Genes of greater or less abundance than Ctrls are labeled in red and blue (P-adj <0.01). (**d**) Gene ontology of all significantly dysregulated genes in c. Top 5 terms are shown if more than 5. (**e**) Progression of the transcriptome defects during meiotic maturation. Genes that are significantly changed at MII stage were selected and sorted according to the $log_2$ FC of *Dis3$^{cKO}$*/Ctrl at GV stage. x axis: transcripts sorted by $log_2$ FC of *Dis3$^{cKO}$*/Ctrl in GV oocytes, and every twentieth transcript was picked to represent the entire transcriptome. (**f**) Violin and boxplot showing distribution of $log_2$ FC of *Dis3$^{cKO}$*/Ctrl at GV, GV3h and MII stages. All transcripts are significantly changed at MII stage. (**g**) Types of genes in each Degradation Potential (DP) group. (**h**) Boxplots showing the change of transcripts of GV3h/GV and MII/GV3h in Ctrl and *Exosc10*$^{cKO}$ oocytes. All transcripts are equally binned into 5 groups by their degradation potential (DP). The number labeled between Ctrl and *Exosc10*$^{cKO}$ in each DP group represents the ratio of the mean of MII/GV3h values in *Exosc10*$^{cKO}$ vs. that in Ctrl.

#### Extended Data Fig. 3

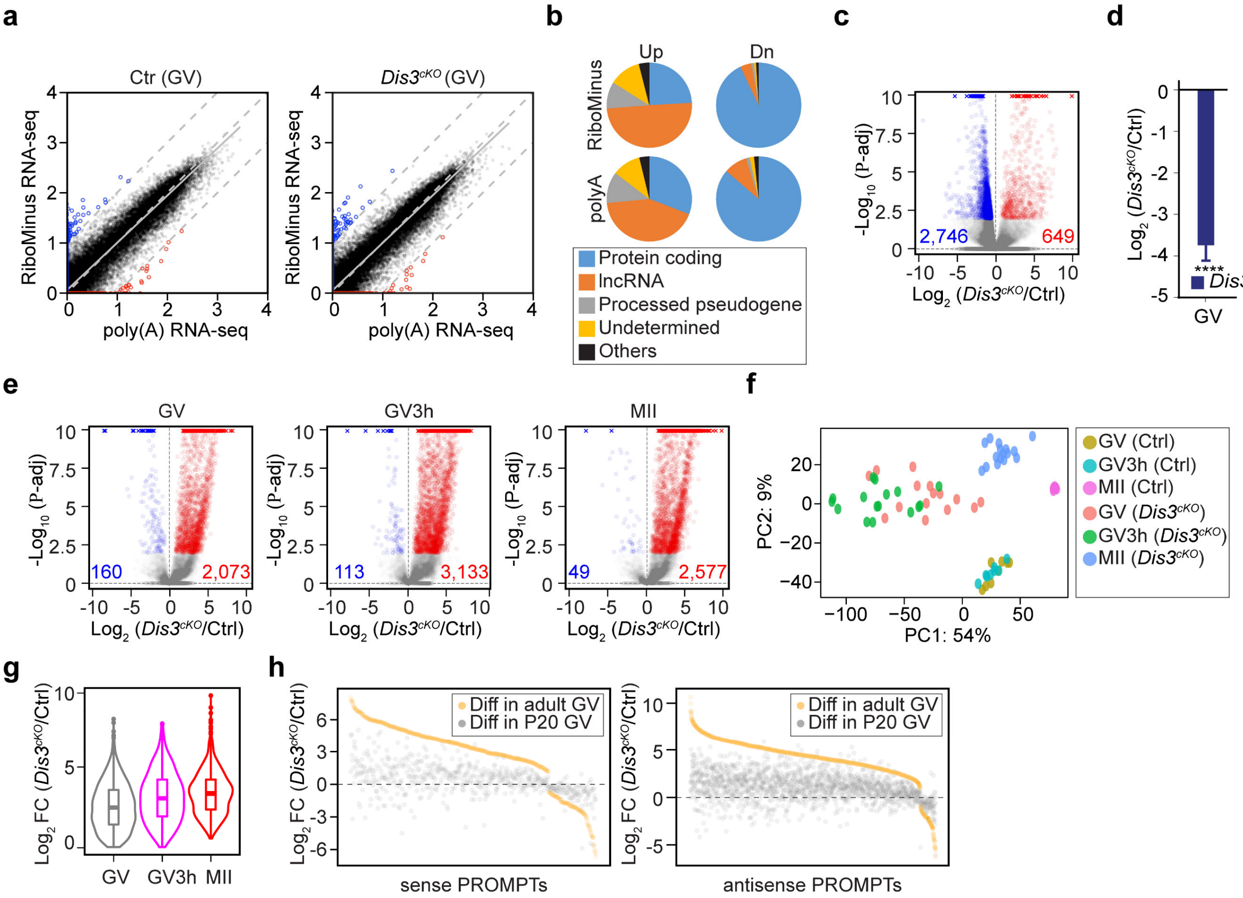


**Extended Figure 3. *Dis3$^{cKO}$* oocytes have accumulated PROMPTs.** (**a**) Correlation plots between RiboMinus RNA-seq and poly (A) RNA-seq of Ctrl and *Dis3$^{cKO}$* GV oocytes. For each gene, x and y are normalized mean of genes of all libraries in each genotype and each method. In each plot, the x-axis is the $\log_{10}$ count value from poly(A) sequencing results and the y-axis is that from RiboMinus sequencing. Red dots, the outliers in poly(A) sequencing; blue dots, the outliers in RiboMinus sequencing. Three gray dashed lines: y=x, y=x-1 and y=x+1. Gray solid line: fitted line of all the dots. (**b**) Types of genes in the up-regulated and down-regulated groups obtained from RiboMinus RNA-seq and poly (A) RNA-seq at GV stage. (**c**) Volcano plot showing changes of genes in *Dis3$^{cKO}$* oocytes at GV stage from RiboMinus RNA-seq. The more and less abundant transcripts are labeled in red and blue (P-adj <0.01). (**d**) Bar graph showing *Dis3* transcript level decreases in *Dis3$^{cKO}$* oocytes at GV stage from RiboMinus RNA-seq. **** P-adj < 0.0001. (**e**) Volcano plots showing changes of PROMPTs abundance, including *Dis3$^{cKO}$*/Ctrl at GV, GV3h and MII stages from poly(A) RNA-seq. The more and less abundant transcripts are labeled in red and blue (P-adj <0.01). (**f**) Dot plot showing PCA of PROMPTs in all samples from poly(A) RNA-seq. (**g**) Distribution of PROMPTs $\log_2$ FC of *Dis3$^{cKO}$*/Ctrl at GV, GV3h and MII stages. (**h**) Accumulation of sense and antisense PROMPTs from P20 stage to GV stage in *Dis3$^{cKO}$* oocytes. x axis: PROMPTs sorted by $\log_2$ FC of *Dis3$^{cKO}$*/Ctrl at GV stage, and every twentieth transcript was picked to represent all PROMPTs RNA. All PROMPTs selected were significantly changed at GV stage.

#### Extended Data Fig. 4

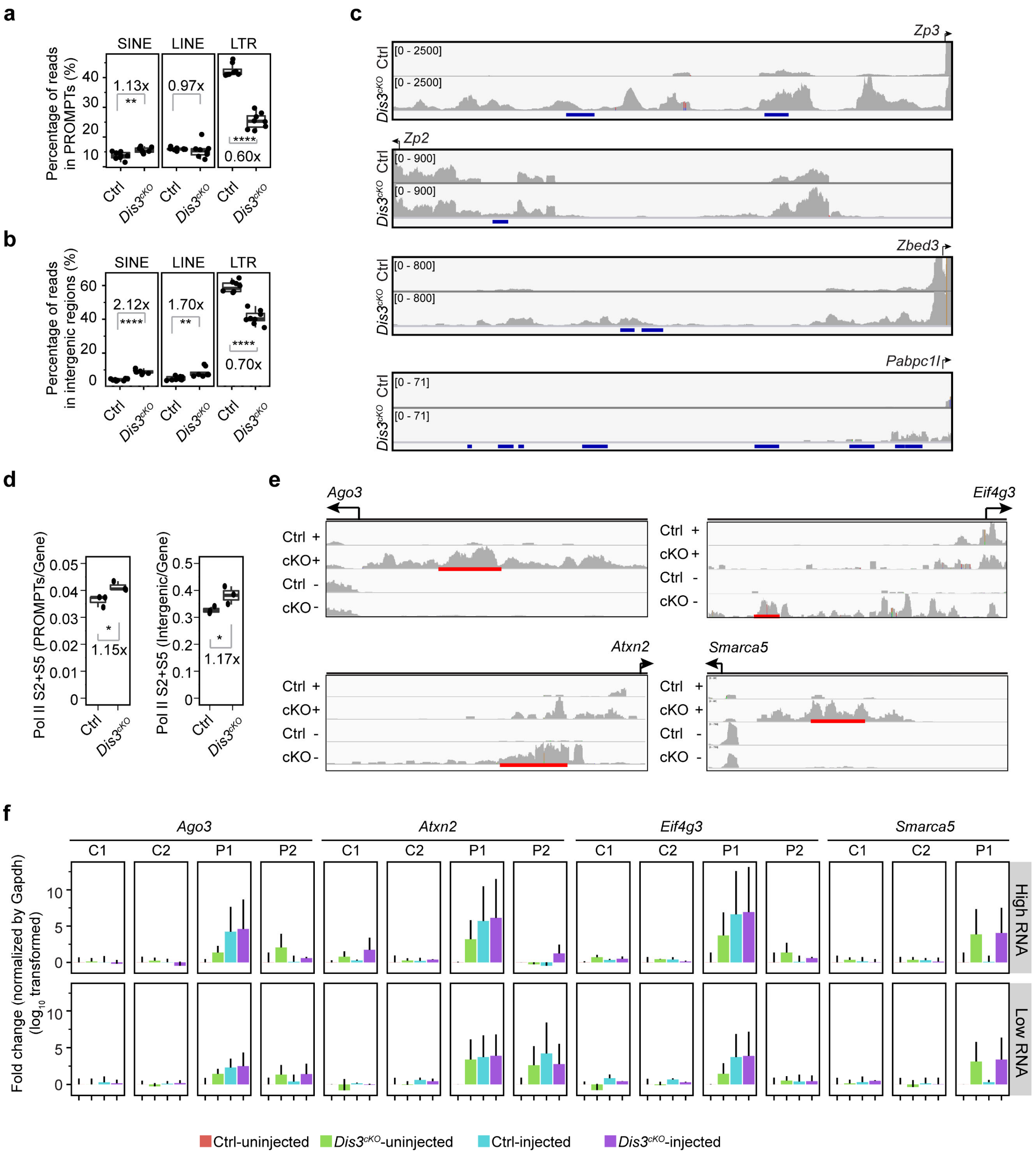


**Extended Figure 4. PROMPTs upregulation associates with SINEs.** (**a-b**) Percentage of reads that were mapped to PROMPTs-located (a) and intergenic (b) SINE/LINE/LTR in Ctrl and *Dis3*$^{cKO}$ oocytes at GV stage. Each dot represents one oocyte sample. The number labeled in each repetitive element panel represents the ratio of the mean percentages of repetitive element-associated reads in *Dis3*$^{cKO}$ oocytes vs. that in Ctrl oocytes. ** P = 0.0053 (PROMPTs-SINE), **** P = 1.11e-09 (PROMPTs-LTR), **** P = 7.13e-08 (intergenic SINE), ** P = 0.0057 (intergenic

LINE), **** P = 1.26e-07 (intergenic LTR). (**c**) Genome browser views showing the PROMPTs regions of *Zp3*, *Zp2*, *Zbed3* and *Pabpc1l*. Dark blue lines represent the SINE loci. (**d**) Pol II S2+S5 occupancy in PROMPTs vs genes, * P =0.03 (left), and in intergenic regions vs genes, * P =0.05 (right). (**e-f**) Microinjection of selected PROMPTs and detection of the microinjected PROMPTs, non-injected PROMPTs and genic region changes by qPCR. Red lines in e indicate the synthesized PROMPT region for microinjection. Note that *Smarca5* PROMPT was used as a control region and was not synthesized for overexpression. All synthesized PROMPTs RNA were mixed for microinjection. Top panel: PROMPTs RNA injected at higher dose (800 ng/μl); bottom panel: PROMPTs RNA injected at lower dose (80 ng/μl). qPCR in f includes the expression level of the microinjected PROMPTs (P1, highlighted in e), non-injected PROMPTs of the same gene (P2), and two genic regions of the same gene (C1 and C2).

#### Extended Data Fig. 5

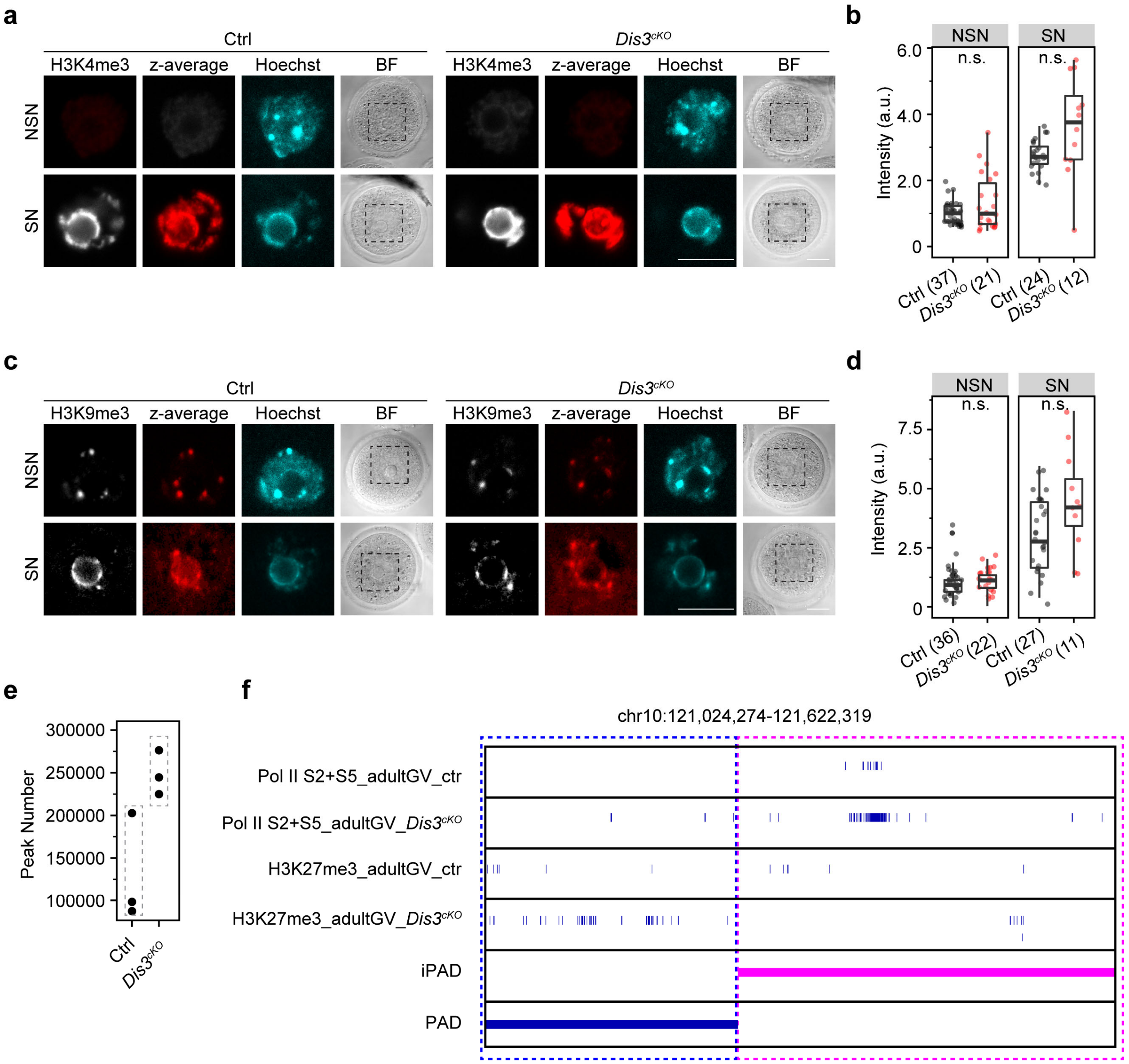


**Extended Figure 5. *Dis3$^{cKO}$* oocytes gain H3K27me3 occupancy in a pre-defined manner.** (**a-d**) Confocal fluorescence and quantification of H3K4me3 and H3K9me3 immunostaining of NSN and SN oocytes at GV stage in Ctrl and *Dis3$^{cKO}$* oocytes. Scale bar: 20 μm. (**e**) Reproducibility of H3K27me3 peaks in Ctrl and *Dis3$^{cKO}$* GV oocytes from CUT&Tag assays. (**f**) An example of genome browser views showing the changes of Pol II S2+S5 and H3K27me3 relative to PAD and iPAD regions.

#### Extended Data Fig. 6

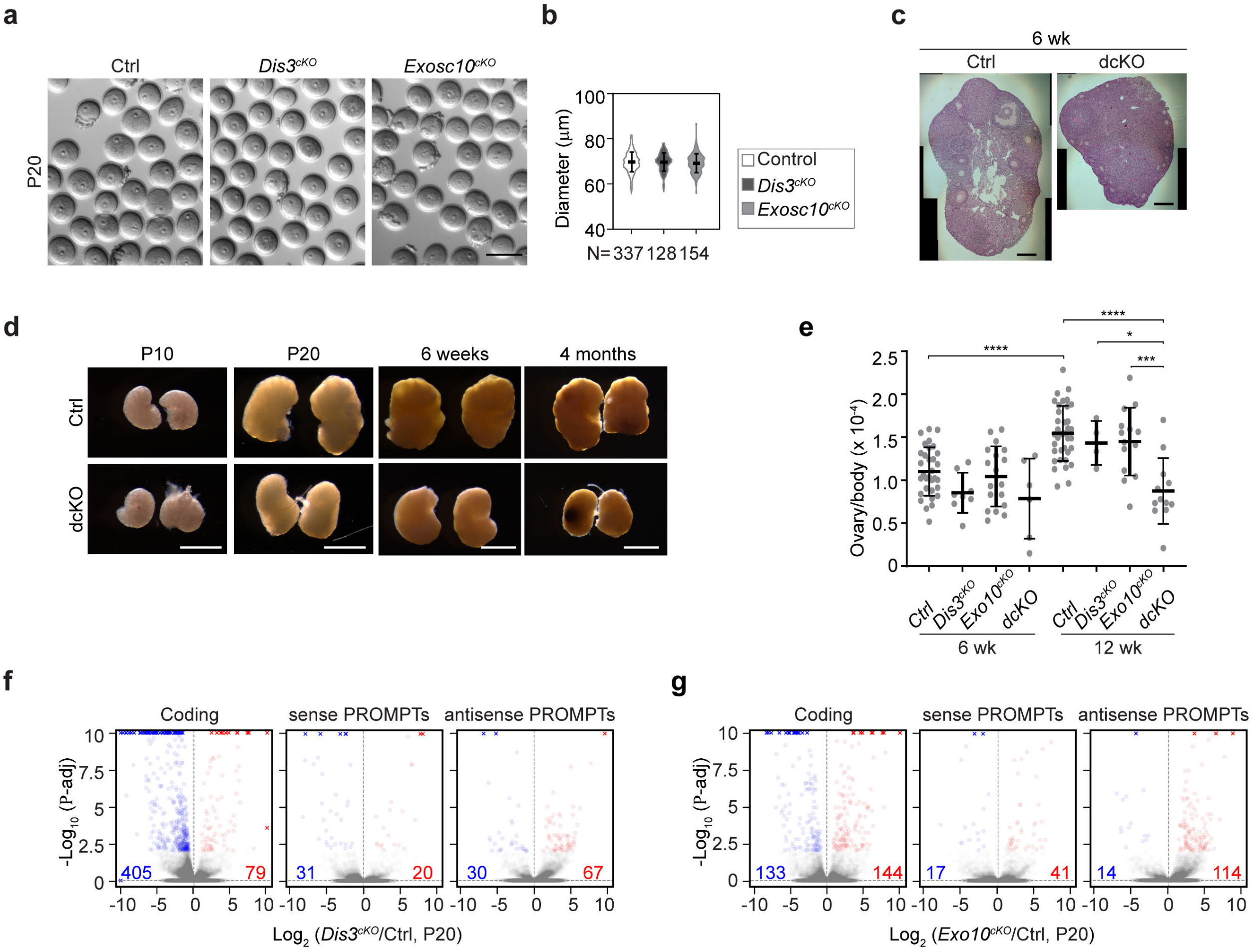


**Extended Figure 6. Oocyte depletion of both DIS3 and EXOSC10 causes growth defects in earlier oogenesis.** (**a**) Live imaging of P20 oocytes in Ctrl, *Dis3*$^{cKO}$ and *Exosc10*$^{cKO}$ females. (**b**) Diameter of oocytes in a. The horizontal lines represent the mean and standard deviation. (**c**) Ovary histology of Ctrl and dcKO females at 6 weeks. Pictures were imaged in tiles and stitched together. (**d**) Ovaries of Ctrl and dcKO females at P10, P20, 6 weeks and 4 months. (**e**) Normalized ovary weight of Ctrl, *Dis3*$^{cKO}$, *Exosc10*$^{cKO}$ and double knockout (dcKO) females at 6 and 12 weeks. **** P = 6.7041E-08 (Ctrl 6 wks vs Ctrl 12 wks), **** P = 4.4387E-05 (Ctrl 12 wks vs dcKO 12 wks), * P = 0.01 (*Dis3*$^{cKO}$ 12 wks vs dcKO 12 wks), *** P = 0.001 (*Exosc10*$^{cKO}$ 12 wks vs dcKO 12 wks) by one-way ANOVA test. The horizontal lines represent the mean and standard deviation. (**f-g**) Volcano plots of genic transcripts (left), sense PROMPTs (middle) and antisense PROMPTs (right) changes in *Dis3*$^{cKO}$ (f) and *Exosc10*$^{cKO}$ (g) oocytes at P20. Scale bar: 100 μm in c, 500 μm in d.

#### Extended Data Fig. 7

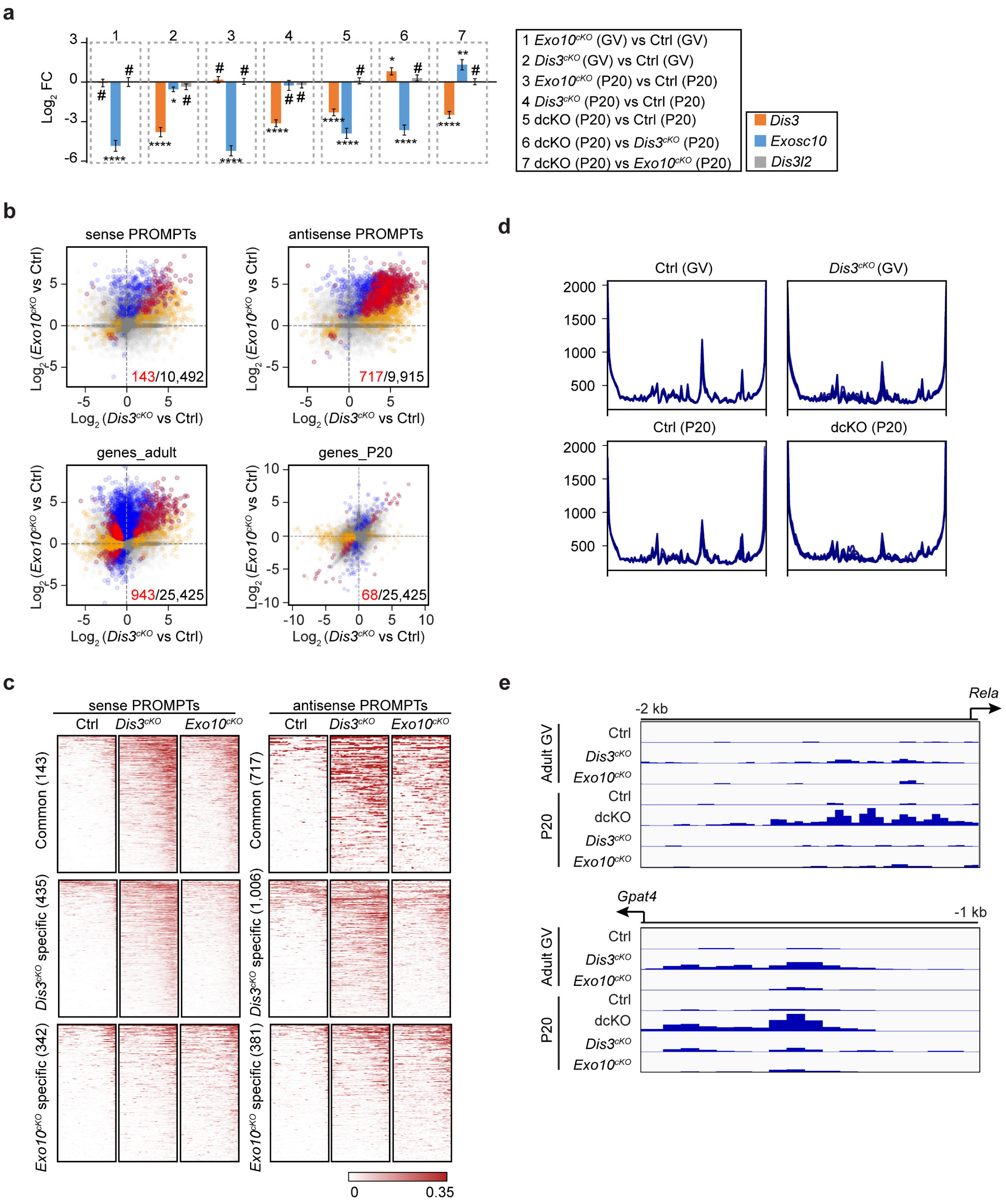


**Extended Figure 7. EXOSC10 facilitates DIS3 in degrading PROMPTs.** (**a**) Bar graphs showing changes of *Dis3* and *Exosc10* transcript levels under different comparisons. Changes of *Dis3l2* transcript level was shown as a reference. (**b**) Top two panels are the correlation plots of differentially expressed sense and antisense PROMPTs in *Dis3*$^{cKO}$ and *Exosc10*$^{cKO}$ GV oocytes. Colors highlighted significantly changed PROMPTs in *Dis3*$^{cKO}$ (orange), *Exosc10*$^{cKO}$ (blue) and both single knockout oocytes (red). Bottom two panels are the correlation plots of differentially expressed genes at adult stage and P20 stage in *Dis3*$^{cKO}$ and *Exosc10*$^{cKO}$ GV oocytes. Color highlighted significantly changed genes in *Dis3*$^{cKO}$ (orange), *Exosc10*$^{cKO}$ (blue) and both single knockout oocytes (red). (**c**) Heatmaps of PROMPTs regions in Ctrl, *Dis3*$^{cKO}$ and *Exosc10*$^{cKO}$ GV oocytes from adult females. PROMPTs were clustered as common, DIS3-specific, and EXOSC10-specific. (**d**) Profiles of RNA-seq read coverage in intergenic regions from Ctrl adult GV oocytes, *Dis3*$^{cKO}$ adult GV oocytes, Ctrl P20 oocytes and dcKO P20 oocytes. Each line represents one oocyte sample. (**e**) Example genome browser views of PROMPTs regions from RNA-seq in different samples.
